# Climatic suitability predictions for the cultivation of macadamia in Malawi using climate change scenarios

**DOI:** 10.1101/2021.05.16.443810

**Authors:** Emmanuel Junior Zuza, Yoseph Negusse Araya, Kadmiel Maseyk, Shonil A Bhagwat, Kauê de Sousa, Andrew Emmott, Will Rawes

## Abstract

Global climate change is altering the suitable areas of crop species worldwide, with cascading effects on people and animals reliant upon those crop species as food sources. Macadamia is one of these essential lucrative crop species that grows in Malawi. Here, we used an ensemble model approach to determine the current distribution of macadamia production areas across Malawi in relation to climate. For future distribution of suitable areas, we used the climate outputs of 17 general circulation models based on two climate change scenarios (RCP 4.5 and RCP 8.5). The precipitation of the driest month and isothermality were the climatic variables that strongly influenced macadamia’s suitability in Malawi. We found that these climatic requirements were fulfilled across many areas in Malawi under the current conditions. Future projections indicated that vast parts of Malawi’s macadamia growing regions will remain suitable for macadamia, amounting to 36,910 km^2^ (39.1%) and 33,511 km^2^ (35.5%) of land based on RCP 4.5 and RCP 8.5, respectively. Alarmingly, suitable areas for macadamia production are predicted to shrink by −18% (17,015 km^2^) and −21.6% (20,414 km^2^) based on RCP 4.5 and RCP 8.5, respectively, with much of the suitability shifting northwards. This means that some currently productive areas will become unproductive in the future, while current unproductive areas will become productive. Notably, suitable areas will increase in Malawi’s central and northern highlands, while the southern region will lose most of its suitable areas. Our study, therefore, shows that there is potential for expanding macadamia production in Malawi. Most, importantly our future projections provide critical evidence on the potential negative impacts of climate change on the suitability of macadamia production in the country. We recommend developing area-specific adaptation strategies to build resilience in the macadamia sector in Malawi under climate change.

## 1. Introduction

Global climate change has become an indisputable fact and has altered ecosystems, including human health, livelihoods, food security, water supply, and economic growth [1]. These impacts are predicted to increase sharply with the degree of warming. For example, warming to 2 °C is expected to increase the number of people exposed to climate-related risks and poverty by up to several hundred million by the 2050s [1]. This represents significant threats to current agricultural production systems in many parts of Africa, especially among smallholder farming families with little adaptive capacity [2], [3]. Sub-Saharan Africa (SSA) is one of the most vulnerable regions to climate change due to the combination of the reduction in precipitation with the increase in temperature [4]–[6]. Within SSA, Malawi has been highlighted as being particularly vulnerable to climate change due to high levels of poverty, limited finances and technology, and heavy reliance on a predominantly rain-fed agricultural sector for its food and nutritional security, economy, and employment [7], [8].

Climate change is already hampering agricultural production in Malawi. Since the 1960s temperatures have increased in all seasons and throughout the country by approximately 0.9 °C [6]. Projected temperature data indicates warming throughout Malawi, from 0.5–1.5 °C by the 2050s. This increase in warming is expected to increase transpiration and evaporation from plants, soil, and water surfaces [9]. Moreover, under all future climate projections (2050–2100), Malawi’s surface temperatures are expected to rise [10]. In terms of precipitation, [6] observed variability in projected amounts and seasonal patterns. Analysis of 34 climate change models projecting up to the 2090s suggests more frequent dry spells and increases in rainfall intensity [11]. These changes are likely to threaten livelihoods, increase the risk of food insecurity, and negatively affect Malawi’s economic growth.

Increased warming and unreliable rainfall within Malawi will impact the landscape and the livelihoods of many rural populations who depend on agricultural activities [10], [12]. Barrueto et al. observed that in Nepal’s highlands, increased temperatures led to changes in cropping systems because of an upward shift in perennial tree crops’ suitability [13]. Like Nepal, climate change will cause shifts in agricultural systems in Malawi. With the country’s high deforestation rates, increases in diurnal temperature changes are expected, which will make it more difficult for crop growth and development [14]. Consequently a proper assessment of climate suitability for various crops under current and future climatic conditions is essential in governing agricultural land use planning in Malawi [12], [15].

Climate suitability reflects the degree of agreement of climate resources required for crop growth and is used to evaluate the relationship between crop distribution and climate factors. Climate suitability, however, only considers climate conditions and does not test socio-economic factors, management practices, and soil types [16]. The process involves applying a crop model to simulate the interaction between different climate factors, explore the potential impact of climate change on agricultural production, and determine production potential for a particular crop and area [17]. Climate suitability assessment should be the first step in agricultural land use planning as it identifies limiting factors for growing a particular crop in an area and aids in decision-making for sustainable agricultural systems [18]. For perennial crops such as coffee and macadamia, climate suitability assessment is essential because they are long-term investments with high initial costs to establish the crops [19]. Proper planning is, therefore, key to the success of perennial tree production. However, in Malawi climate suitability studies have primarily focused on important cash and staple crops [20]. Cash crops are cultivated mainly to be sold rather than used by the people who grew them [12]. Important cash crops in Malawi include tea [21], cashew, coffee, cotton, pulses, sugarcane, tobacco [12]), and macadamia [20]. Cash crops such as macadamia and pulses contribute to food security and economic development, particularly among the producers as they are used for income generation. Staple crops are consumed routinely and in large quantities and constitute a dominant portion of the standard diet. In Malawi, the most important staple crop is maize [20].

Agriculture forms the backbone of the economy and society of Malawi [22]. Nearly 85% of the country’s households are dependent on agricultural activities for their livelihoods [23]. The agricultural sector consists of two distinct sub-sectors: smallholder farmers and commercial estates sub-sectors. About 11% of the rural labor force works on commercial estates to supplement farm income, and around 80% is engaged in the smallholder sub-sector [24]. Smallholder production accounts for 90% of all the country’s food [25]. Despite the smallholder sub-sector contributions to Malawi’s food security, individually, most smallholders are food insecure annually [26]. This is due to their dependency on rainfed agriculture, limited usage of modern tools for farming, and unpredictable weather patterns [26], [27]. Moreover, food security among these smallholder farmers is not permanent, as the fall from food abundance to food scarcity can occur within a matter of days when one’s income is lost to bad weather [28], [29]. Meeting Malawi’s growing demand for food in the coming decades is likely to become more difficult as already stressed agricultural systems will be challenged by population growth (expected to peak at 38.1 million by 2050) and rising incomes, especially among rural communities. Thus, effective agricultural adaptation to the changing climate conditions requires a good understanding of how climate change may affect cultivation patterns and various crops’ suitability.

Studies on the projected effects of climate change in Malawi have mainly focused on staple and cash crops, specifically maize and tobacco. Little is known about perennial tree crops, which are essential to addressing the country’s future food security uncertainty [30], [31]. However, a good understanding of how climate change may affect cultivation patterns and crops’ suitability is an effective agricultural adaptation strategy for survival. This is because adverse effects of climate change in the future may drive smallholder farmers and rural populations, in general, to migrate due to a reduction in productivity and changes in land-use zones. To mitigate the projected impacts of climate change on land and crop use planning, Benson et al. examined the climate suitability of a wide range of important crops grown in Malawi [12]. But, this study did not evaluate the suitability of macadamia in Malawi despite its increasing importance as a commercial cash crop and its benefits for food and nutrition security.

Macadamia (*Macadamia integrifolia* Maiden & Betche) is an evergreen perennial crop and belongs to the Proteaceae family [32]. Its kernel contains more than 72% oil content and is one of the most highly regarded nuts globally. This is due to its high nutritional value and high market price driven by consumers’ high demand for the nuts and products [32]. Macadamia trees were historically inter-cropped with coffee, tea, and tung oil in commercial estates in southern Malawi [33]. Currently, over one million macadamia trees have been planted under the commercial estate sub-sector and over 300,000 trees under the smallholder sub-sector, which is expected to increase to over 1,000,000 in the next decade [20]. Globally, Malawi is the seventh-largest producer of macadamia nuts [20]. According to [20], [33], Malawi could become the biggest producer of macadamia in the world. This potential is attributed to the country having the most suitable altitude and climatic conditions for its growth and development and large land pockets among smallholders that offer expansion opportunities.

Macadamia is a lucrative crop among smallholder producers in Malawi for food security and income generation [34]. However, climate change is expected to negatively affect productivity and land suitability for macadamia production in the future in most producing countries. This is because macadamia is sensitive to variations of climatic factors, especially cannot resist higher temperatures and droughts [35]. Macadamia is best suited in areas with annual mean temperatures (Tmean) ranging from 10–15 °C [36]. The optimal temperature for macadamia growth and development is between 16–30 °C [36]. Nevertheless, lower day temperatures (≤10 °C) are lethal to the crop. Nagao and Ho found that higher temperatures exceeding 30 °C are associated with water loss that subsequently restricts the build-up of oil in the nuts and reduces their quality [37]. For precipitation, various authors have suggested a tolerable annual rainfall ranging from 510 to 4000 mm [35–38]. Britz reported that macadamia yield in South Africa was lost due to spatiotemporal variability in precipitation and temperature [39]. A detailed summary regarding climatic conditions for macadamia production is given in Supplementary Table S1.

A scientific description of climate suitability of crop distribution is of great significance to mitigate the negative effects of climate change and ensure food security. Therefore, this research aims to assess the suitability of macadamia in Malawi’s current and future climate and predict suitable geographic regions for its production. First, we assess the current spatial distribution of macadamia in Malawi. Then we model the future distribution of macadamia utilizing bioclimatic variables [40], obtained from downscaled Coupled Model Intercomparison Project 5 (CMIP5) GCMs [41], based on two emission scenarios (RCP) of climate change [42]. We focus on climate projections for the 2050s to align with the United Nations framework of global challenges in agriculture and food security [43].

## 2. Methods

### 2.1. Study area

Malawi falls within the longitudes 30 and 40, and the latitudes −17 and −10 (Supplementary Fig S1). The country spans over 118, 484 km^2^, with 94, 449 km^2^ (80%) of land area and 24, 035 km^2^ (20%) of water surfaces. Soil nutrient status varies tremendously across the country due to the variability in topography (Fig 1), parent materials, and management, especially among smallholder farmers [15]. Due to this reason, soil characteristics were not considered for our analysis.

**Fig 1:**
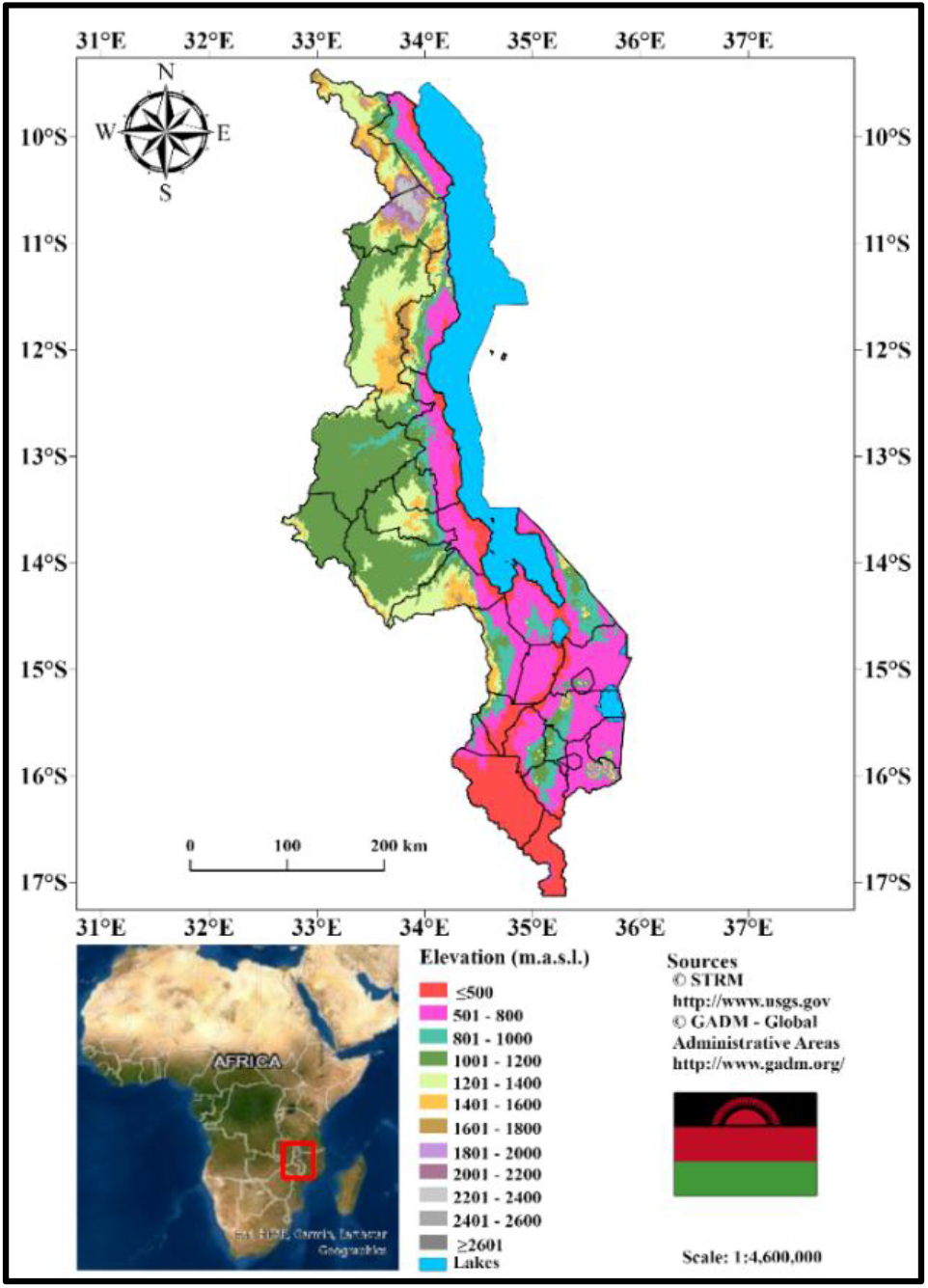
Geographic location and topography of Malawi based on Shuttle Radar Topography Mission digital elevation model data.

Malawi has a sub-tropical climate with two distinct seasons, the rainy season from November to April that delivers 90–95% of the annual precipitation [44], and the pronounced dry season from May to October. The geographical distribution of temperature and precipitation in Malawi is primarily determined by topography and the distance to the Indian Ocean and Lake Malawi. Further, large elevation changes in escarpment areas make the climate in the uplands and lowlands significantly different [44]. Lower maximum temperatures and higher precipitations are experienced in different escarpment areas in Malawi. For example, the northern parts of Malomo in Ntchisi district lies in the rain shadow area, while the southern part receives higher rainfalls and cooler temperatures. The average precipitation varies from 500 mm in low-lying marginal areas (≤ 500 meters above sea level/ m.a.s.l.) to over 3000 mm in high plateau areas [12]. The mean annual minimum and maximum temperatures for Malawi are 12 and 32 °C, respectively, with the lowest temperatures in June and July and the highest in October or early November [45]. Fig 2 illustrates the spatial pattern of average annual temperatures (a) and annual precipitation (b).

**Fig 2:**
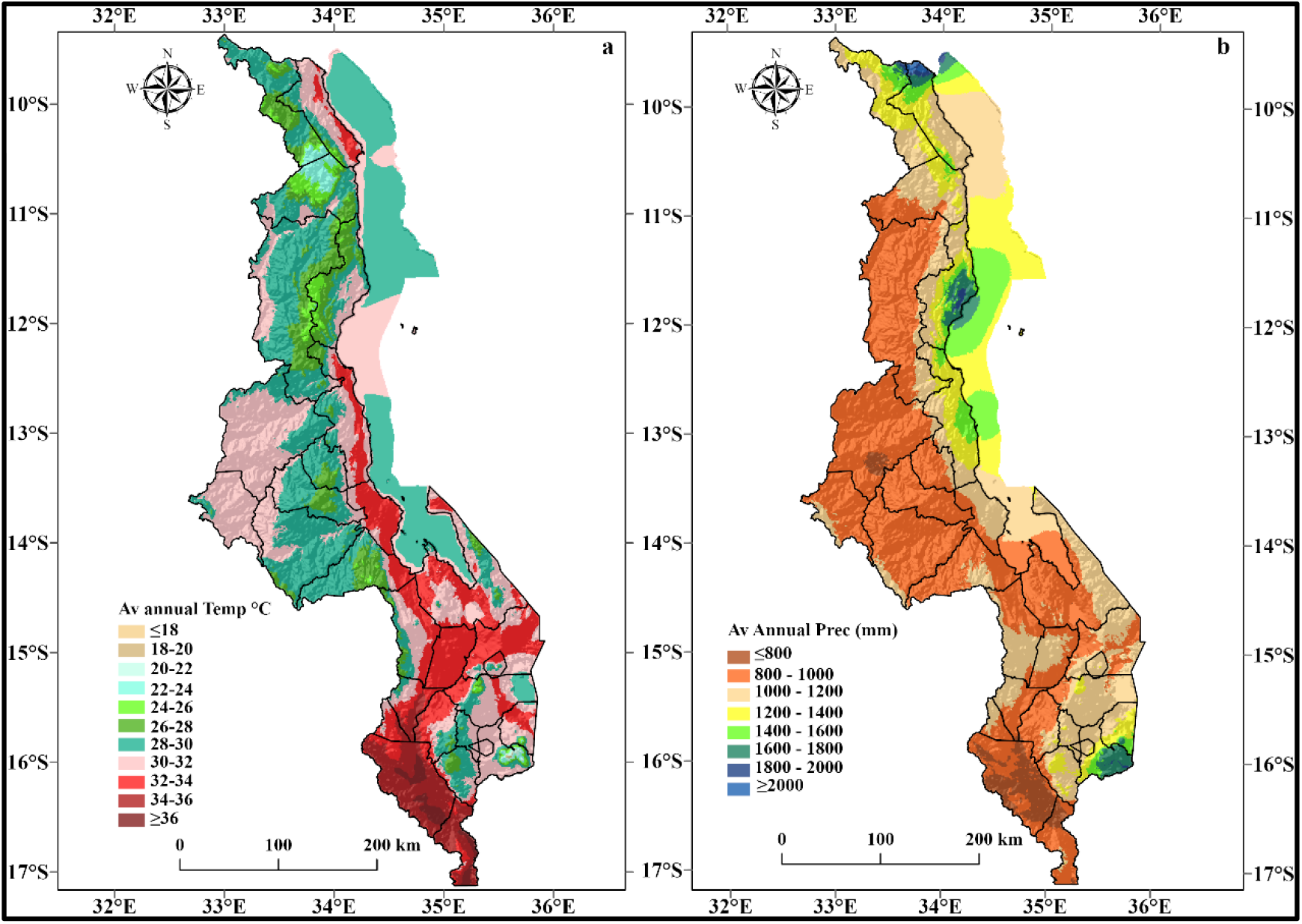
a). Average annual temperature (°C) and b). Precipitation (mm) of Malawi based on WorldClim-Global Climate Data.

### 2.2. Occurrence data

Data on macadamia tree species’ occurrence was collected from smallholder macadamia producing districts in 2019 through a field survey of macadamia farms in Malawi. For our analysis, we only sampled ten-year-old successfully established macadamia orchards under smallholder rainfed conditions. At each farm, the Global Position System (GPS) coordinates (in WGS84 datum) were collected using a global position system (Garmin eTrex Vista® Cx) together with altitude. A total of 120 orchards were sampled throughout Malawi, but for this study, a total of 36 points were used for the modelling purpose (Fig 3). The remaining 84 occurrence points were used for cross-validation to evaluate the predictive model accuracy [46]–[49].

**Fig 3:**
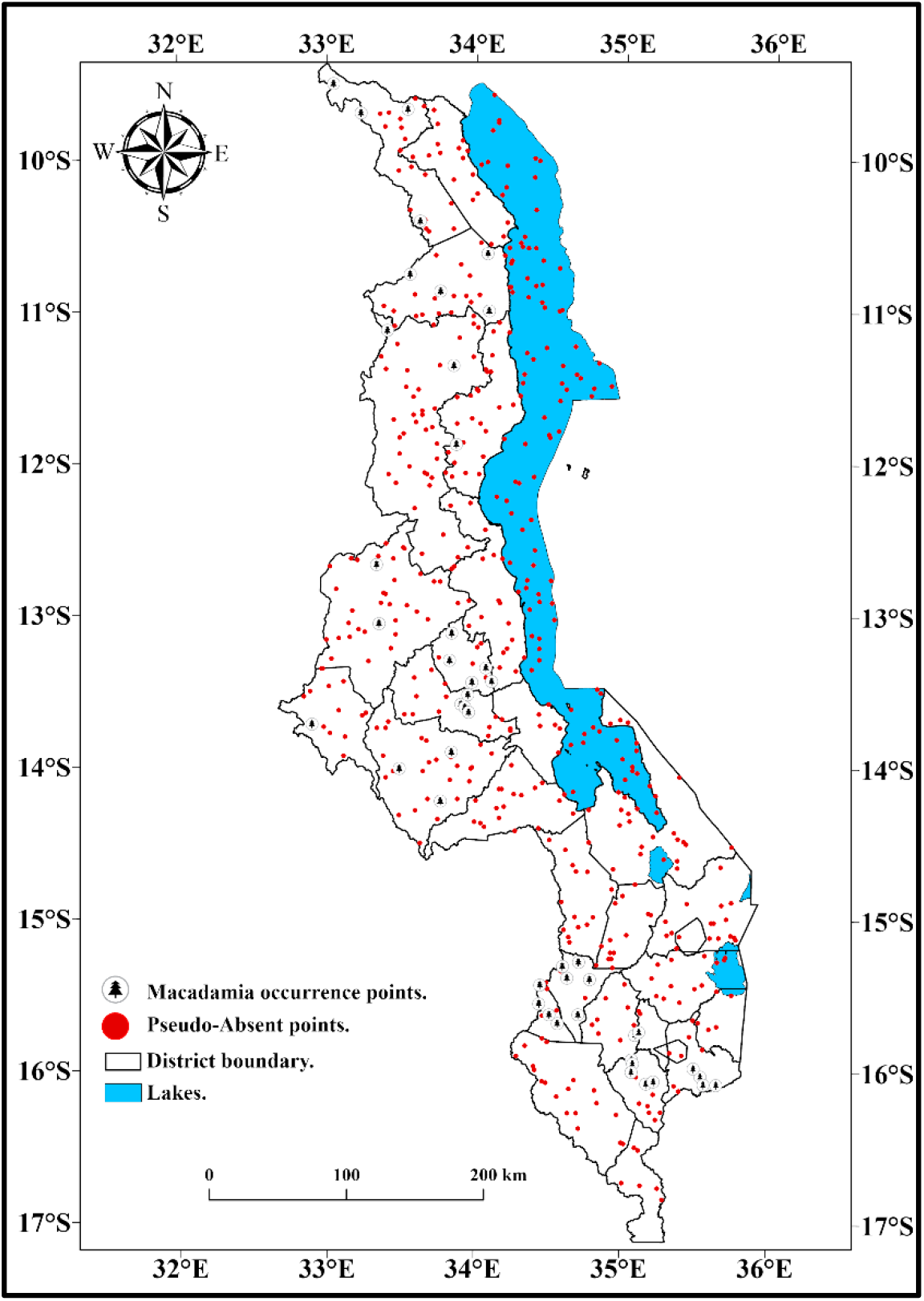
Map of Malawi showing macadamia occurrence points and pseudo absent points.

### 2.3. Climate data

We used bioclimatic predictors (~1970–2000) from WorldClim data set version 1.4 (http://www.worldclim.org/) at a spatial resolution of 2.5 arc-minute (4.5 km^2^ at the equator) to model the current areas suitable for macadamia production in Malawi. Calculated from monthly temperature and precipitation climatologies, these variables reflect spatial variations in annual means, seasonality, and extreme/limiting conditions (Supplementary Table S2). We used bioclimatic variables from 17 general circulation models (GCMs) based on two representative concentration pathways (RCP) of climate change [42] for future predictions. We selected RCP 4.5, which is an optimistic scenario that considers an intermediate GHG concentration and predicts an average increase in temperature by 1.4 °C (0.9–2.0 °C) and RCP 8.5 the most pessimistic scenario, which considers higher GHG emissions concentration with a 1.4–2.6 °C projected increase in mean global temperature by the 2050s (period 2046–2065). The bioclimatic variables from WorldClim that we used include limiting factors that are ecologically important based on temperature and precipitation variation. To avoid model overfitting, we selected the least correlated variables by applying the variance inflation factors (VIF) and retained those with VIF < 10 [50]. Variables with the highest correlation (VIF ≥ 10) were removed, resulting in eight bioclimatic predictors for our analysis (**Error! Reference source not found.**). The long-term ecological conditions are essential for predicting perennial crop production [51] because perennial crops are in the field for more than 25 years, and productivity is measured by yield quantity and quality.

### 2.4. Modelling approach

We modelled the current and future distribution of macadamia species in Malawi based on an ensemble suitability method implemented by the R package BiodiversityR. We used an ensemble modelling technique because it combines predictions from different algorithms and can provide better accuracy in predictions than relying on individual species distribution models [52]. The procedure consisted of four steps.

Firstly, we evaluated the predictive performance accuracy of 18 algorithms of species distributions models (SDM) using a fivefold cross-validation technique. Following work by [53] and [54], we divided the occurrence data into two different sets by randomly assigning 70% of the data as a training dataset to fit the model, and the remaining 30% were used as test data to evaluate the model prediction accuracy. To test the stability of the prediction accuracy, a five-fold (bins) cross-validation replicate was performed in the model as described by [47]. Each SDM algorithm’s performance was evaluated from each bin separately after individual algorithms were assessed with data from the other four bins. The algorithms’ performance was measured with the area under the curve (AUC). The AUC value provides a specific measure of model performance, demonstrating the model’s ability to locate a randomly selected present observation in a cell of higher probability than a randomly chosen absence observation [52], [55]. We used an AUC value of 0.77 as a threshold to select the best-performing algorithms for our analysis. Species distribution model algorithms that did not fit this criterion were not used to calculate the ensemble model’s suitability [56].

Additionally, we only used SDM algorithms that can distinguish between suitable and non-suitable areas without needing absence locations [57]. The presence-only approach was utilized because, for agricultural applications of niche models, it is inappropriate to treat areas without current production as entirely unsuitable. Further, it is difficult and rare to determine whether a species is absent in a particular location. Hence, absence data may not represent naturally occurring phenomena [51]. According to [58], presence-only models can produce reliable predictions from limited presence datasets, meaning that they are robust and a cheaper option for obtaining training datasets. To enhance our ensemble model’s predictive ability, the macadamia occurrence data was coupled with 500 randomly pseudo-absence data generated throughout Malawi (Fig 3). We used the pseudo-absence data as opposed to using real absences to avoid underestimation issues [59].

In the second step, we retained only the algorithms that contributed at least 5% to the ensemble suitability (*Se*) [50]. This generated AUC values for each algorithm and the parameters of the response functions that estimated the probability values of species occurrence based on the climate of each grid cell of the study area (Supplementary Table S5). AUC values ranged between 0 and 1, and a value less than 0.5 indicated that the simulated result was worse than random [60]. Classification of model performance estimated by the AUC was: 0.50–0.60 fail; 0.60–0.70 poor; 0.70–0.80 fair; 0.80–0.90 good; 0.90–1.0 excellent, and further the AUC value higher than 0.77 [61]. We later combined the results of all the models by calculating for each model the weighted average (weighted by AUC for each model) of each model’s probability values to generate the ensemble suitability map. We weighted the AUC values using the following equation:

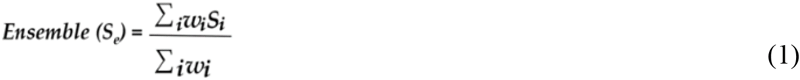

Where the ensemble suitability (*Se*) is obtained as a weighted (*w*) average of suitabilities predicted by the contributing algorithm (*S_i_*).

The third step generated the current distribution maps (probability maps and presence-absence maps) of macadamia under the current climate. This was based on the weights which were generated during model calibration. To generate the absence-presence layers, we used the maximum sensitivity (true positive^+^) and maximum specificity (true negative^-^) approach [62], where we reclassified the distribution maps to binary maps (suitable and unsuitable areas). In [48], [59], [63], it was shown that this method is one of the most reliable for choosing a reclassification method. In this analysis, sensitivity is the proportion of observed presences correctly predicted and therefore is a measure of omission errors, whereas specificity represents the proportion of correctly predicted absences and thus quantifies commission errors.

To create distribution maps for future bioclimatic conditions, we utilized the same procedure used in the baseline suitability and presence-absence maps but utilized the climate information from each of the 17 future GCM for RCP 4.5 and RCP 8.5. Since no criteria exist to assess which of the GCMs best predict future climate [64], by incorporating all 17 GCMs, we encompassed all possible changes in the distribution of the macadamia species. To integrate the results of the 17 GCMs presence-absence layers into a single layer, we used the criterion of likelihood scale [48], which requires at least 66% of agreement among GCMs to keep the predicted presence or absence in a given grid cell.

## 3. Results

### 3.1. Factors determining land suitability of macadamia in Malawi

Our study has shown that precipitation-related variables were the most important in determining the distribution and suitability of macadamia in Malawi. Precipitation of the driest month (9.69) was the variable with the greatest relative influence on macadamia production. Possibly because of the sensitivity of pod growth during this phase to water scarcity. Among the temperature variables, isothermality (this variable is calculated by dividing mean diurnal temperature range by mean annual temperature range) was the most significant, with a VIF score of 8.95 (Table 1). Based on our ensemble model, annual means did not influence macadamia suitability in Malawi.

**Table 1:** Climate variables influencing macadamia suitability in Malawi.

| Covariate | Bioclimatic variable | Unit | VIF Score |
| --- | --- | --- | --- |
| Bio14 | Precipitation of driest month | mm | 9.69 |
| Bio3 | Isothermality (BIO2/BIO7) x 100 | - | 8.95 |
| Bio15 | Precipitation seasonality (cv x 100) |  | 8.09 |
| Bio2 | Mean diurnal range (Mean of monthly) | °C | 8.05 |
| Bio18 | Precipitation of warmest quarter | mm | 7.01 |
| Bio13 | Precipitation of wettest month | mm | 6.73 |
| Bio6 | Minimum temperature of the coldest month | °C | 5.87 |
| Bio4 | Temperature seasonality (Std. Dev x 100) | - | 4.12 |

### 3.2. Current suitability of macadamia in Malawi

Results of the present (~1970–2000) suitability analysis showed that 53,925 km^2^ (57.4%) of the surface area in Malawi is suitable for macadamia production (Fig 4), while 40,524 km^2^ (42.6%) is unsuitable for the crop. Therefore, our findings demonstrate that currently, macadamia is grown in a broad range of environments (Fig 4); the variations in the color gradient represent the degree of macadamia suitability per grid cell. Suitability values range from 0 and 1, whereas values of ≥ 0.84 (forest green) are considered as highly suitable areas, 0.74–8.4 (jade green) optimal, 0.54–0.74 (mint green) moderate, and ≤ 0.54 (yellow) indicate marginal areas. We observe that macadamia’s optimal suitability was across the higher elevated areas and marginal suitability in the lower elevated areas. Further, suitability peaked in areas around 1000 m.a.s.l., receiving at least 1000–1200 mm of rainfall and temperatures not exceeding 30 °C.

**Fig 4:**
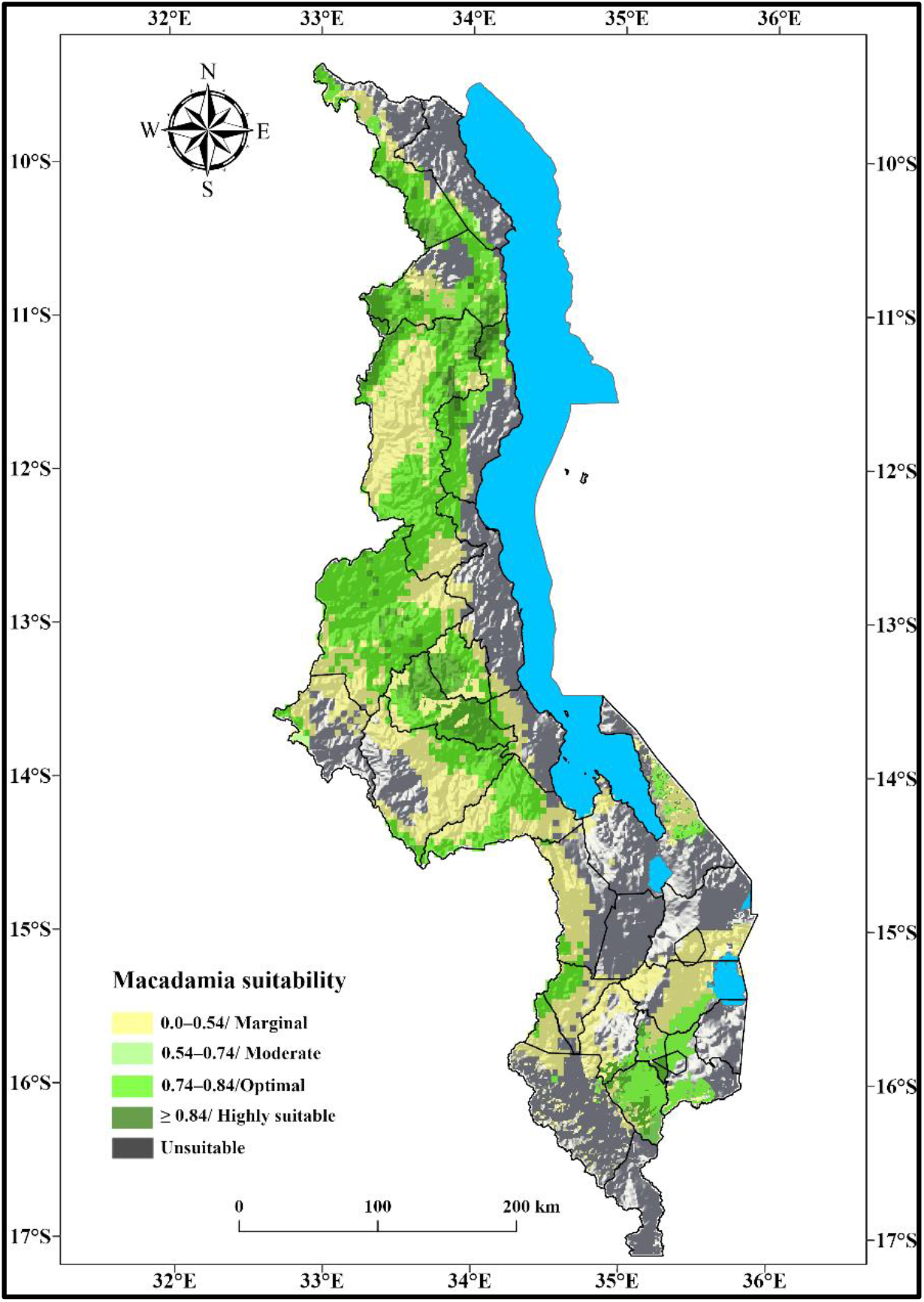
Current suitability for macadamia production in Malawi.

Interestingly, our findings showed optimal suitability in some areas where the average annual temperatures are considered too hot for macadamia (≥ 30 °C), specifically in areas around Katunga (Chikwawa), Luchenza (Mulanje), and Nsabwe (Thyolo). Highly suitable areas were observed in Malawi’s mountainous regions with elevation ranging from 1200–1600 m.a.s.l. in some parts of Dowa, Chitipa, Machinga Mulanje, Mzimba, Ntchisi, Nkhatabay, Rumphi, and Thyolo. Areas with moderate suitability were predicted in the mid-hills between 950–1000 m.a.s.l. in Blantyre, Chiradzulu, Dedza, Kasungu, Lilongwe, Mchinji, Mwanza, Neno, Ntcheu, and Zomba districts. Marginally suitable areas are found in the lower elevated (≤ 900 m.a.s.l) parts of Malawi. Expectedly, our ensemble model for the current distribution of suitable areas for macadamia production largely overlapped the area of macadamia production in Malawi. Additionally, these areas are also utilized for the production of other crops, especially annuals.

### 3.3. Gain and loss of suitability under future projections in Malawi

Compared to current climate conditions, the extent of suitable areas for macadamia production is expected to decrease in the future under both emission scenarios utilized in this analysis. Our results revealed a net loss of *-*18% and *-*21.6% of potentially suitable land for macadamia production under RCP 4.5 and RCP 8.5, respectively (Fig 5). This translates to 17,015 km^2^ (RCP 4.5) and 20,414 km^2^ (RCP 8.5) of Malawi’s total cultivatable surface area. Areas located in lower altitudes (500–1000 m.a.s.l.) will suffer the greatest decline in suitability due to the projected general temperature increases and reduced precipitation amounts and distribution. These losses will be more pronounced in Malawi’s southern region areas, especially those along the shire valley. Further, some southern region areas will become marginal or even unsuitable for macadamia, while others will remain suitable though less than today. Thyolo district, which is currently the country’s most productive and biggest macadamia growing area, is predicted to suffer significant reductions in suitability areas due to climate change. This is attributed to southern Malawi’s low-lying nature and high risks of heatwaves, flooding, and droughts linked to the El Niño Southern Oscillation.

**Fig 5:**
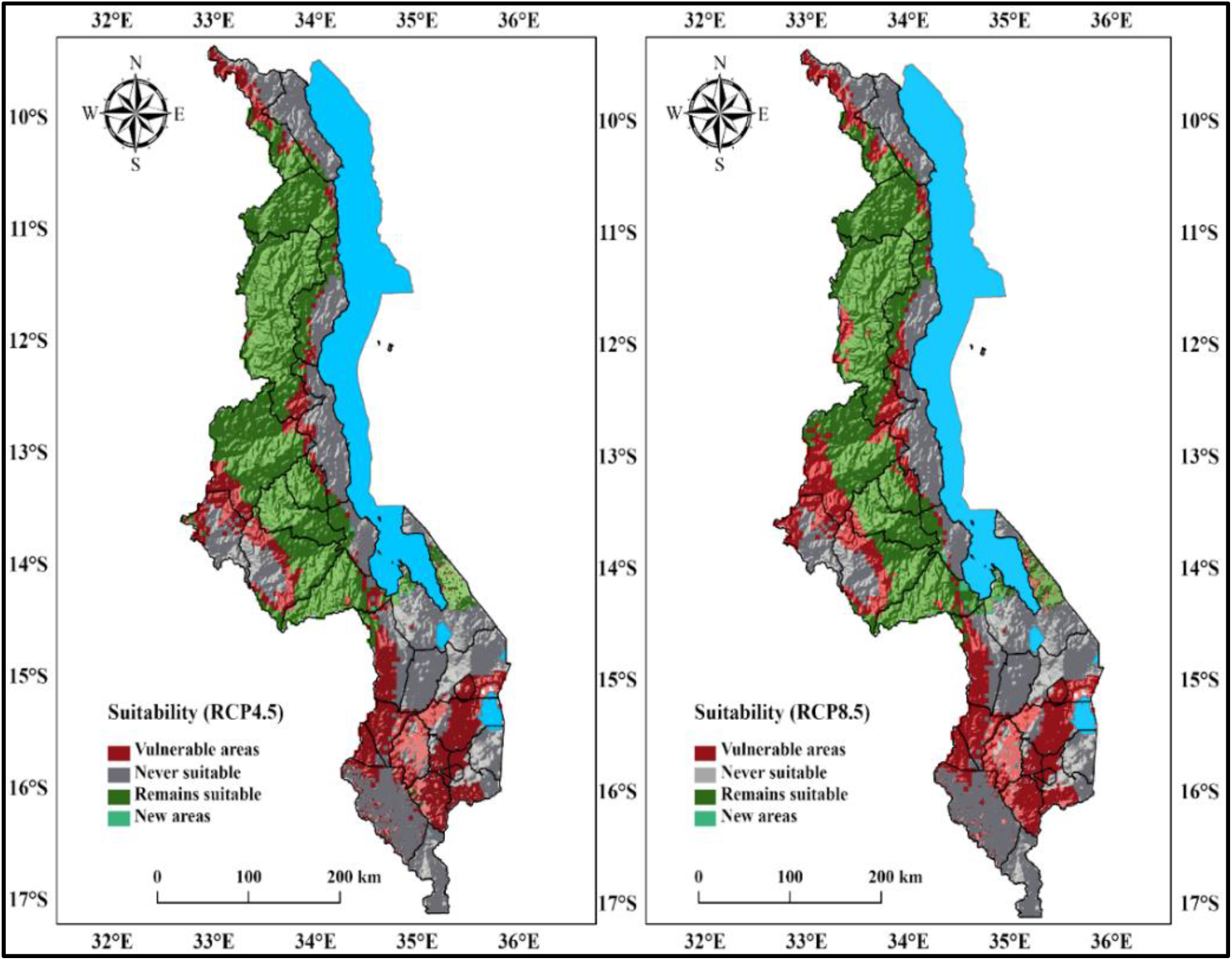
Shifts in macadamia suitability due to climate change by 2050 (a) RCP 4.5 (b) RCP 8.5.

Surface gains under future climate projections for macadamia suitability are described in Table 2. Both scenarios show that a large fraction of suitable areas for macadamia production will remain unchanged. Approximately 36,910 km^2^ and 33,511 km^2^ of Malawi’s surface area are projected to remain unchanged under RCP 4.5 and RCP 8.5, respectively. The intermediate optimistic scenario (RCP 4.5) indicated an average gain (newer areas) of 0.22% of Malawi’s surface area, amounting to approximately 207 km^2^ of potentially suitable land. Under RCP 8.5 scenario, newer areas are projected to account for 0.5% of the land, translating to 476 km^2^ of Malawi’s total surface area. We observed that projected newer areas will be more under RCP 8.5, amounting to 0.28% more than RCP 4.5. The reason being that some of the very cold areas currently unsuitable for macadamia will become suitable due to the projected increased warming by the scenario RCP 8.5. The newer areas are predicted to occur in Dedza (Mua and Chipansi), Mangochi (Namwera and Chaponda), and Ntcheu (Tsangano and Bonga) districts based on both emission scenarios. Nevertheless, these apply only to very limited areas in the country and cannot compensate for the suitability decrease in the lowlands. Our analysis, therefore, shows that the results for the RCP 4.5 and RCP 8.5 models are similar in direction, but the RCP 8.5 models project a greater reduction in suitable areas in warmer locations and expansion of suitable areas in colder locations by the 2050s.

**Table 2:** Future distribution area of macadamia production in Malawi by 2050.

| RCP | Category | Area (km<sup>2</sup>) | % changes<sup>a</sup> | Net change (%)<sup>b</sup> |
| --- | --- | --- | --- | --- |
| 4.5 | No longer suitable | 17,015 | -18.0 |  |
|  | Never suitable | 40,317 | 42.7 |  |
|  | Remains suitable | 36,910 | 39.1 | -17.8 |
|  | New Areas | 207 | 0.22 |  |
| 8.5 | No longer suitable | 20,414 | -21.6 |  |
|  | Never suitable | 40,047 | 42.4 |  |
|  | Remains suitable | 33,511 | 35.5 | -21.1 |
|  | New Areas | 476 | 0.5 |  |

<sup>a</sup> Percentage of the total land area of Malawi (94,449 km<sup>2</sup>).
<sup>b</sup> Net change is the balance between colonization and loss (positive net balance indicates an increase in the areas suitable for the species, and negative net balance indicates a decrease in the areas suitable for the species).

Our results suggest a northward shift in the location of the most suitable areas for macadamia production, a reduction of highly suitable areas in the south, and an increase along the central and northern parts of Malawi, dependent on the landscape topography (Fig 6). Areas projected to lose their suitability occur mainly in Malawi’s southern regions, including Blantyre, Chikwawa, Chiradzulu, Machinga, Mwanza, Mulanje, Thyolo, and Zomba districts. The projected loss is approximately 95–100% of the currently suitable areas in southern Malawi. This is attributed to the projected increases in temperature and frequency of droughts in the areas. Nevertheless, some higher elevated areas (≥ 1600 m.a.s.l.) within Chitipa (Misuku hills), Ntchisi (Malomo and Kalira), and Rumphi (Mphompha and Ntchenachena) districts will similarly lose some of their suitable areas for macadamia production.

**Fig 6:**
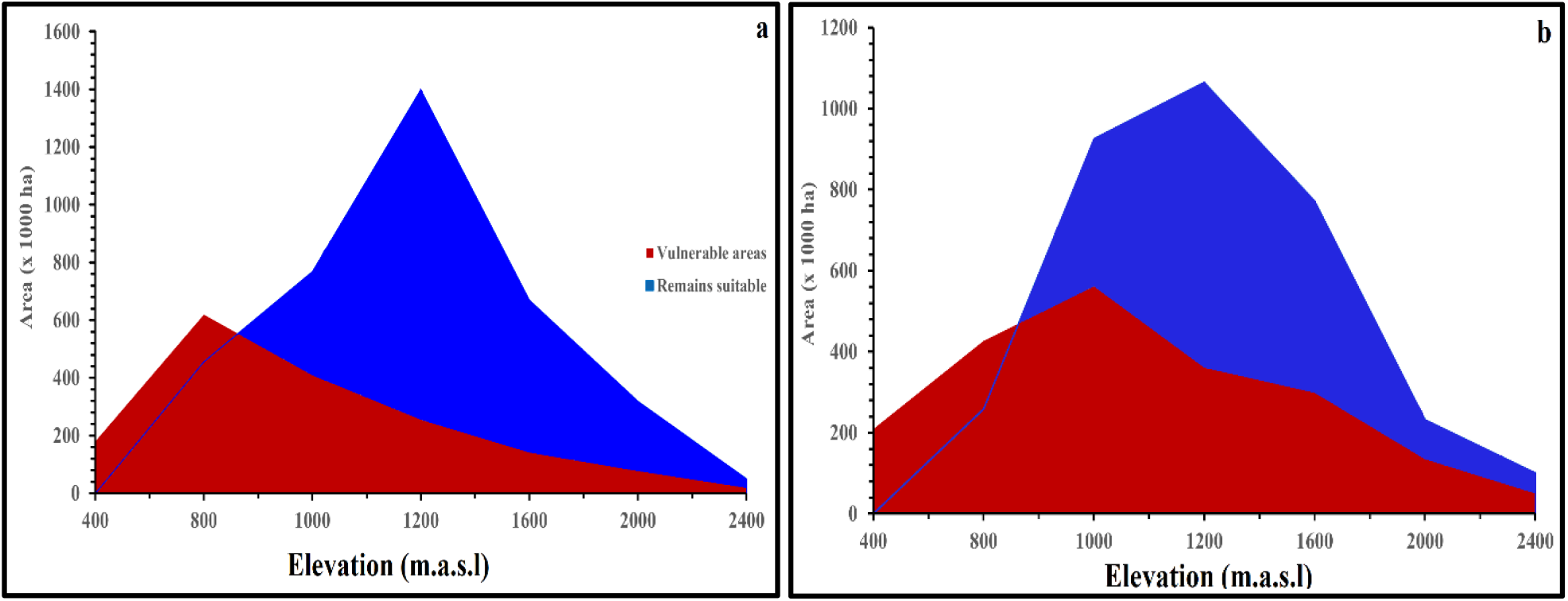
Shifts in macadamia suitability by 2050 in Malawi shown by the altitudinal gradient (a) RCP 4.5 (b) RCP 8.5.

## 4. Discussion

Precipitation and temperature have been identified as critical factors influencing crop growth and yields across the globe. Our results revealed that macadamia suitability in Malawi is influenced by the interactions of temperature and precipitation and seasonal variations of the two than the annual means. The climate variables at a national scale determined in this study might be different from climate indicators at the regional and global scale. A previous researcher showed that the annual mean temperature, the warmest month’s maximum temperature, minimum temperature of the coldest month, and annual precipitation were viewed as climatic regionalization indices for macadamia in Nepal. In the present study, we observe that precipitation-based factors are more valuable in determining the suitability of macadamia in Malawi, verifying zoning studies for macadamia production done for the country [38]. In these studies, precipitation distribution and quantity were identified as the most critical variables, and it is apparent when considering the current macadamia belt that water supply is the most limiting factor for the crop. Consequently, projections that climate change will reduce rainfall amounts making its distribution unreliable in many parts of Malawi, especially the southern region [9], will drive many areas out of macadamia production.

The distribution of precipitation is more related to precipitation of the warmest quarter and precipitation seasonality. In this regard, areas with sufficient and sustainable water supply during the drier months of the season (May–November) will remain suitable for macadamia. However, the regions with low annual precipitation and soils with poor water holding will lose their macadamia production suitability. Our findings concur with early studies by [65], who found that water deficits from prolonged drought periods induced macadamia flower losses and tree mortality, leading to lower yields in Australia. An interview with one of the macadamia farmers in Malomo in Ntchisi district identified the link between water scarcity and macadamia suitability. Joseph Makono explained in the interview:

> *“During the flowering period which coincides with the dry season, I have observed that most of the macadamia flowers drop from the trees because of the low moisture content in the soil around the plant.”*

To avoid yield losses caused by drought stress, farmers need to adopt moisture conservation measures (mulching, rainwater harvesting, box ridging, and basins) and possibly develop irrigation infrastructure to match the water requirements for macadamia growth and development annually. This is especially important for the areas in southern Malawi that are prone to droughts and flooding.

Temperature isothermality was found to be the most important factor determining the suitability of macadamia in Malawi. This index is a measure of temperature heterogeneity and is a composite of two variables reflecting temporal variation in temperature: diurnal range and annual range. Our findings indicated that large fluctuations between day and night temperatures and increased warming affect macadamia suitability in Malawi. Marginal suitability of the crop was observed in areas located in the hotter (≥ 30 °C) and lower elevated parts of Malawi, notably along the lakeshore and Shire valley. This is attributed to higher daytime and night-time temperatures experienced in these areas. Optimal suitability is observed in intermediate to upper elevated areas that experience cooler temperatures, especially at night. Thus, projections that climate change will increase the number of hot days (30.5) and hot nights (40) [11] will certainly reduce the number of suitable areas for macadamia production in Malawi due to increased warming, which will result in increased evapotranspiration rates. Taking this into account, trees currently grown in the hotter areas will require sufficient water availability to cater to the water lost through evapotranspiration. In Australia, Nepal, and South Africa, studies have shown that high daytime and high night-time temperatures are responsible for the reduction in yields and suitable production areas for macadamia [36], [39], [66], therefore agreeing with our current findings in Malawi. Consequently, climate change will have dual impacts on macadamia production by reducing suitable production areas and reducing the nut yield and quality.

Due to its geographic location and socio-economic status, Malawi is most exposed to climate change [7], [8]. Thus understanding species’ response to climate change is crucial in Malawi for agricultural land use planning, notably for high-value perennial crops [44], [49], [56]. Our predictions suggested that extensive areas in Malawi under the current climatic conditions are suitable for macadamia production. At present, the crop is grown over a wide range of altitudes (500–1400 m.a.s.l.) throughout the country. Furthermore, our findings suggested the suitability of macadamia in Malawi’s south-eastern parts, such as in Luchenza, Katunga, and Nsabwe, which are beyond the current reported production areas and considered to be too hot for the crop. This is expected as the suitability maps capture the potential production areas, some of which have not yet been translated to realized areas [51]. Additionally, this illustrates the broad adaptability of some macadamia cultivars that allows its production from high potential areas to marginal and low input areas with several environmental constraints. Nonetheless, these areas are the most vulnerable to climate change because of limited buffering potential.

Malawi is already falling outside the prescribed optimal range for macadamia production, attributing it to climate change. This is evident by the 0.9 °C increase in annual mean temperature and overall drying recorded in the past five decades [6], [67], [68]. As a result of the projected temperature increases, changing rainfall patterns, and increased water scarcity, the suitability for macadamia production in Malawi is likely to decrease in the 2050s and is expected to shift northwards. Differences in loss-gain of suitability highlight which agro-ecological zones could be more vulnerable to climate change (Fig 7). According to our predictions, lowland areas will be the most affected (due to inadequate rainfall), with the central and northern highlands even improving capacity to sustain macadamia production in areas where this is not possible due to environmental constraints. Other authors have predicted similar shifts in the suitability of macadamia-producing areas caused by the impacts of climate change. Barrueto *et al.* reported an upward shift in suitable areas for macadamia production in Nepal due to the negative impacts of climate warming [13]. Platts *et al.* found that in the United Kingdom, species will shift their distributions polewards and to higher elevations in response to climate change [69]. Being associated with a particular set of environmental conditions, it is feared that Malawi may lose some of its suitable areas for macadamia production due to climate change. Consequently, our findings highlight the negative impacts climate change may have on macadamia suitability in Malawi.

From our results, we observe that the extent of suitable areas for macadamia production in Malawi will decrease over the next 40–50 years. Our analysis reveals that the currently suitable areas in the southern region will be the most affected, while areas located along the country’s central and northern parts, dominated by highlands, will become more favorable for the crop. The greatest victims will be areas currently experiencing a hotter and drier environment (Fig 8). Consequently, these results show the sensitivity of macadamia to variations in ecological conditions. Our findings confirm and, more importantly, extend the work by [38], who found an inverse relationship between increases in temperature (all the four RCPs) with the decline in suitability for macadamia production in Nepal. Other published studies show that higher temperatures (≥ 30 °C) and water stress reduce macadamia vegetative growth and reproduction [70], restrict the build-up of oil [71], reduce raceme and nut retention [72], and reduce macadamia yields [73]. Therefore, in areas where there are no predicted macadamia suitability changes, farmers could continue planting their macadamia trees. However, both research and field-based evidence from discussions with farmers show that climate-related changes are already occurring (heatwaves, droughts, and flooding) and affecting macadamia production in Malawi. For example, Kelvin Masinga, a macadamia farmer from Neno district, reported that:

> *“Recently, I have started seeing the effects of climate change on my macadamia trees, the coats of the nuts have changed their color from dark green to brown, and due to very hot weather, there is increased failing and dying of flowers.*”

Farmers are, therefore, encouraged to start implementing adaptation measures such as the use of improved macadamia varieties, agroforestry, intercropping, water conservation, and irrigation for long-term and sustainable macadamia production. Nonetheless, these suitability changes are predicted to occur over the next 40–50 years, so these will mostly impact the next generation of macadamia farmers rather than the current generation. Therefore, there is still time for adaptation. Failure to adapt in time to the risk of decreasing yields and incomes may lead to the migration of rural populations to the main cities of Blantyre, Lilongwe, and Mzuzu.

Altitude provides an excellent climatic change comparison for health, growth, and yield of crops [74], [75]. As a result, individual plants grow very well in high altitudes, whereas others can only grow in middle or lower-altitude areas [76]. Comparing the current and future suitable areas for macadamia production in Malawi reveals an upslope shift in suitability. Our ensemble model showed that low-lying areas at altitudes ranging from 500–1000 m.a.s.l. will have a decline in macadamia suitability because of the projected general temperature increases and more dryer conditions. This is primarily true to Malawi’s southern districts, mainly those along the lake and shire valley (Blantyre, Mwanza, Neno, Mulanje, Chikwawa, Thyolo, and Zomba). This is in line with the predicted losses in land suitable for tea production in the same region (Mulanje and Thyolo) due to projected increases in warming and droughts [21]. Similarly, in their global study of coffee suitability, [76] reported that climate change might lead to large losses of areas suitable for the coffee across the globe, mostly in low altitudes below 1000 m.a.sl. However, we established that some higher elevated areas (≥ 1600 m.a.s.l.), such as some parts of Chitipa, Nkhatabay, Ntchisi, and Rumphi, will lose suitability due to predicted cold temperatures (≤ 4 °C) and frequent and intense rainfall (≥ 1750 mm). This reduced suitability is attributed to the high levels of cloud cover experienced in these areas, which results in lower light intensity reaching the leaves of the trees, thus affecting the total net photosynthesis for tree growth and oil accumulation. Our findings coincide with [77], who found that suitable areas for macadamia production decreased after an increase in altitude of over ≥ 1400 m.a.s.l in Thailand. Despite large areas losing suitability, our findings show that some areas will gain suitability for growing macadamia. This will generally depend on the landscape topography and will occur in the mid-altitude areas as suitability moves upslope to compensate for increased temperature. Nevertheless, this only applies to minimal areas within Malawi and cannot compensate for the decrease in suitability.

### 4.1. Applicability and potential limitations of this study

Species distribution modelling in space and time is founded on assumptions intrinsic in the models, some of which cannot be tested [46], [78]. Although this study’s findings can be considered robust, several issues should be considered in the interpretation and application of the results. Though we identified areas as suitable for macadamia production based on environmental factors, however on the ground, this may not directly translate to the size of the arable land. Other physical and socio-economic (including the gender and age of the smallholder farmers, availability of agricultural advisory services, and market availability) factors that determine suitability are difficult to capture in this type of modelling and should be considered in applying model results. Due to these challenges, the authenticity of models in making predictions is questioned [76–77], but modeling remains an important tool for future planning purposes [60], [78], [79]. Therefore, the need for a thorough evaluation of adaptation approaches suggested for smallholder macadamia farmers, as these may be different from those utilized by commercial growers.

It is known that SDM development, particularly for areas with varying topographical terrains such as that of Malawi, is challenging due to the complexity of the local and regional climate gradients [82]. Hence careful interpretation is required when utilizing our results for the local effects of the future predictions on macadamia production in Malawi. For agricultural land use planning, our results must be interpreted with the knowledge of soil nutrition and social-economic factors. In addition, not all macadamia cultivars may be similarly affected by climate change. We recommend that further studies need to be conducted to evaluate the effect of climate change on the trait combination of the various cultivars available in Malawi. This will ensure that the right cultivar is grown in the right place to maximize yields.

The temperature and precipitation data utilized in our analysis are based on the IPCC Special Report on Emission Scenarios [83] using CMIP5 model ensembles (RCPs). We used both emission scenarios and model ensembles from the IPCC Firth Assessment Report (AR5) for our modeling analysis. However, our analysis did not consider Malawi’s economic conditions provided by the Shared Socioeconomic Pathways (SPPs). The SPPs provides five distinct narratives (where each SPPs aligns with one or two of the RCPs) about the future of the world, exploring a wide range of plausible trajectories of population growth, urbanization, economic growth, technological and trade development, and implementation of environmental policies [58], [84]. Therefore a combination of RCPs and SSPs in a model provides more distinct future scenarios that are more feasible due to the integration of radioactive forcing (W/m^2^) and socio-economic development influences [83], [86]. Despite the lack of incorporating the SSPs in our modelling approach, our results’ interpretation and recommendations combine the ensemble model’s prediction results, our general knowledge of Malawi’s climate and agricultural systems, and expert opinions. Therefore, our study is considered thorough in providing accurate and meaningful results for macadamia’s current and future suitability.

Model building is another limiting factor when considering to assess the distribution of species in an area due to different forms of uncertainties that may be incurred during this process [79]. We utilized the automated model calibration method for our analysis as it is embedded with novel modelling frameworks [79]. Using the automated approach, we eliminated sources of uncertainty such as collinearity and model overfitting, which are associated with other methods of model building, such as that of the “priori selection of a set of explanatory variables” model building method [79], [87]. Consequently, our results have a high accuracy level (AUC, 0.88) because we reduced uncertainty caused by highly correlated environmental predictors by applying the variance inflation factors during variable selection. In addition, unlike smallholder farmers, agricultural planners are required to take a long-term view of the situation and rely on predictions from models to support their decisions. The cost of being wrong can be very high. As such, the idea of using ensemble models for suitability studies is vital and appealing [48], [55], [88], [82]. This is because ensemble models combine predictions from different algorithms of SDM and the results are closest to the truth in all circumstances [17], [90], [91].

## 5. Conclusions

In responding to climate change impacts, the United Nations Framework Convention on Climate Change (UNFCCC) advocates for Least Developed Countries’ response to be adaptation rather than mitigation. Therefore, building a sustainable and climate-resilient macadamia sector in Malawi could provide a much-needed economic boost. Our ensemble model successfully delineated the current climatically suitable areas for macadamia production and the potential expansion of suitable areas by the 2050s. However, most of the suitable areas identified for macadamia production exist in agricultural land currently utilized for the production of other crops. Therefore, we suggest promoting macadamia agroforestry and intercropping with other crops such as maize, groundnuts, soybeans, and sunflower in the agricultural fields as an adaptation strategy to climate change farm intensification due to limited land, particularly among smallholders. Additionally, large mono-cropped orchards are generally at high risks for pests and disease. However, these risks would be kept minimal if macadamia is planted in small-scale agroforestry plots, which is what we recommend.

Future projections have indicated northward shifts in areas suitable for macadamia production in Malawi. Extensive areas currently suitable for the crop are projected to be lost in the future due to increased warming and extreme precipitation patterns, especially droughts. Nevertheless, some new areas will become suitable for the crop. Therefore, macadamia communities need to develop locally specific adaption measures for the macadamia sector’s continued profitability under adverse climatic conditions for increased resilience. Among the priority measures to reduce Malawi’s macadamia sector’s vulnerability to climate change is breeding programs for greater drought resistance. To be effective, the varieties and traits need to be selected with the inclusion of smallholder farmer preferences. Irrigation might be an option in some places; however, considering its cost is unlikely to be adopted by a large number of smallholder farmers in Malawi.

We conclude that further research should examine other factors that influence macadamia production within Malawi’s agro-ecological zones to improve our ensemble species distribution model’s applicability. For example, vital research on market accessibility, availability of agricultural advisory services, workload, and gender perspectives among macadamia smallholders should be conducted for informed decision-making. On a national level, we recommend research into the influence of expected climate changes on microclimatic and soil conditions such as soil texture, soil pH, soil nutrition, wind, and humidity. We also recommend investing in studies that examine and employ better quality techniques of planning, selecting, and cultivating the best crop varieties for Malawi’s climate. Finally, at a strategic level, we strongly recommend an investigation into the impact of climate change on the current land use policy and its implications for agriculture in Malawi, especially for macadamia and other high-value perennial crops that require significant initial investments.

## Supporting information

https://zenodo.org/record/4751439#.YJw-o7VKhEZ

## Acknowledgements

The authors are grateful to the Open University and the UK Research and Innovation through Global Challenges Research Fund (GCRF) project for funding and academic guidance. Thanks go to Dr. Phillip Holden, The Open University, UK, Prof. Rick Brandenburg, North Carolina State University, USA, Dr. Abel Chemura, Potsdam Institute for Climate Impact Research (PIK), German, Dr. Michael G. Chipeta, Oxford University, Dr. Edith B. Milanzi, MRC Clinical Trials, University College London, the Neno Macadamia Trust, UK and Highlands Macadamia Cooperative Union Limited (HIMACUL) smallholder farmers, Malawi. We further express our gratitude to Mr. Ken Mkangala and Nicholas Evans for their constructive comments and feedback on the state of macadamia production in Malawi. However, mistakes and omissions are our responsibility.

## Author contributions

**Conceptualization:** Emmanuel Junior Zuza, Yoseph N. Araya, Kadmiel Maseyk, Shonil Bhagwat, Andrew Emmott.

**Data curation :** Emmanuel Junior Zuza, Kauê de Sousa.

**Formal analysis :** Emmanuel Junior Zuza, Kauê de Sousa.

**Methodology :** Emmanuel Junior Zuza, Kauê de Sousa, Kadmiel Maseyk, Yoseph Araya, Shonil Bhagwat.

**Software :** Emmanuel Junior Zuza, Kauê de Sousa.

**Supervision :** Yoseph N. Araya, Kadmiel Maseyk, Shonil Bhagwat, Andrew Emmott.

**Validation :** Emmanuel Junior Zuza, Yoseph N. Araya, Kauê de Sousa, Kadmiel Maseyk, Shonil Bhagwat, Andrew Emmott.

**Visualisation :** Emmanuel Junior Zuza, Kauê de Sousa, Yoseph N. Araya, Shonil Bhagwat.

**Writing – original draft :** Emmanuel Junior Zuza, Yoseph Araya, Kadmiel Maseyk, Shonil Bhagwat.

**Writing – Review and Editing :** Emmanuel Junior Zuza, Kauê de Sousa, Kadmiel Maseyk, Shonil Bhagwat, Yoseph N. Araya, Andrew Emmott.

