## Supplementary material for "Climatic suitability predictions for the cultivation of macadamia in Malawi using climate change scenarios": https://zenodo.org/record/4751439#.YJw-o7VKhEZ

**This PDF file includes:**

Supplementary text

Figs. S1 to S2

Tables S1 to S6

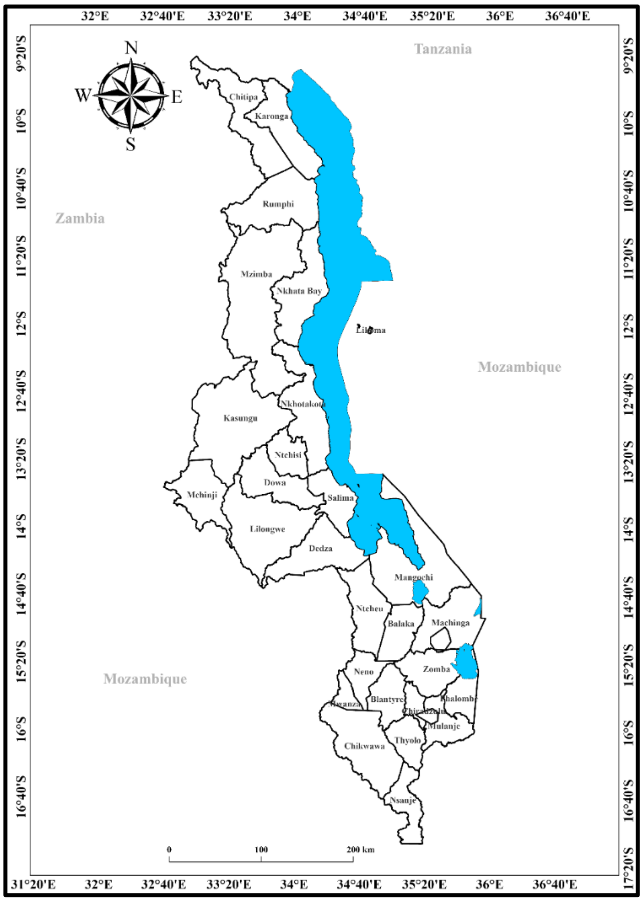


**Fig. S1.** The geographic location of Malawi.

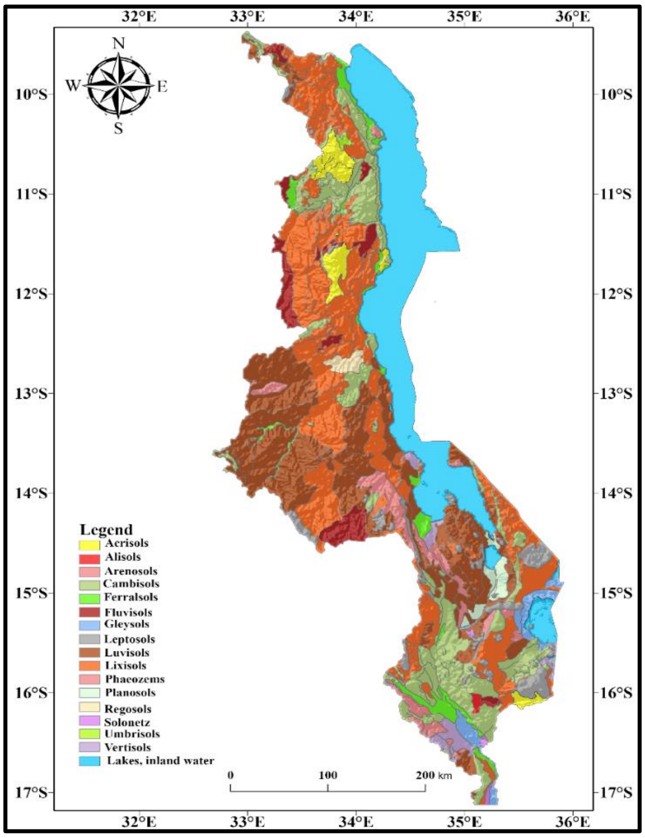


**Fig. S2.** Soil types in Malawi.

**Taxonomy of macadamia**

**Table S1**: Suitable climatic conditions for macadamia production in Malawi.

| Description | Category | Adverse | Moderate | Optimal |
|---|---|---|---|---|
| Minimum temperature of the coldest month. | $T_{min}$[ºC] | ≤1 | 1–4 | 5–10 |
| Annual mean temperature. | $T_{mean}$[ºC] | ≤9 | 10–15 | 16–30 |
| Maximum temperature of the warmest month. | $T_{max}$[ºC] | ≥36 | 31–35 | 25–30 |
| Annual precipitation. | Prec[mm] | 0–700 & ≥1750 | 900–1000 & 1300–1750 | 1000–1250 |

**Table S2**: Bioclimatic variables available in WorldClim.

| Covariate | Bioclimatic variable | Unit |
|---|---|---|
| Bio1 | Annual Mean Temperature | ºC |
| Bio2 | Mean Diurnal Range (Mean of monthly) | ºC |
| Bio3 | Isothermality (BIO2/BIO7) x 100 | - |
| Bio4 | Temperature Seasonality (Std. Dev x 100) | - |
| Bio5 | Max Temperature of Warmest Month | ºC |
| Bio6 | Min Temperature of Coldest Month | ºC |
| Bio7 | Temperature Annual Range | ºC |
| Bio8 | Mean Temperature of Wettest Quarter | ºC |
| Bio9 | Mean Temperature of Driest Quarter | ºC |
| Bio10 | Mean Temperature of Warmest Quarter | ºC |
| Bio11 | Mean Temperature of Coldest Quarter | ºC |
| Bio12 | Annual Precipitation | mm |
| Bio13 | Precipitation of Wettest Month | mm |
| Bio14 | Precipitation of Driest Month | mm |
| Bio15 | Precipitation Seasonality (cv x 100) | - |
| Bio16 | Precipitation of Wettest Quarter | mm |
| Bio17 | Precipitation of Driest Quarter | mm |
| Bio18 | Precipitation of Warmest Quarter | mm |
| Bio19 | Precipitation of Coldest Quarter | mm |

**Table S3**. The general circulation model (GCM) used to obtain climatic variables under scenarios RCP 4.5 and RCP 8.5 in 2050.

| Country | Modelling centre | GCM | Abbreviation |
|---|---|---|---|
| Australia | Commonwealth Scientific and Industrial Research Organization | ACCESS1-0 AC | AC |
| China | Beijing Climate Center | BCC-CSM1-1 BC | BC |
| USA | National Center for Atmospheric Research | CCSM4 CC | CC |
| France | Centre National de Recherches Météorologiques, Centre Européen de Recherche et de Formation Avancée en Calcul Scientifique | CNRM-CM5 CN | CN |
| USA | Geophysical Fluid Dynamics Laboratory | GFDL-CM3 GF | GF |
| USA | NASA/GISS (Goddard Institute for Space Studies) | GISS-E2-R GS | GS |
| South Korea | National Institute of Meteorological Research, Korea Meteorological Administration | HadGEM2-AO HD | HD |
| | | HadGEM2-CC HG | HG |
| UK | Met Office Hadley Centre | HadGEM2-ES HE | HE |
| Russia | Russian Academy of Sciences, Institute of Numerical Mathematics | INMCM4 IN | IN |
| France | Institut Pierre-Simon Laplace | IPSL-CM5A-LR IP | IP |
| | Atmosphere and Ocean Research Institute (The University of Tokyo), National Institute for | MIROC-ESM-CHEM MI | MI |
| Japan | Environmental Studies, and Japan Agency for Marine-Earth | MIROC-ESM MR | MR |
| | Science and Technology | MIROC5 MC | MC |
| Germany | Max Planck Institute for Meteorology | MPI-ESM-LR MP | MP |
| Japan | Meteorological Research Institute | MRI-CGCM3 MG | MG |
| Norway | Bjerknes Centre for Climate Research, Norwegian Meteorological Institute | NorESM1-M | NO |

**Table S4.** Characteristics of climate change scenarios by the 2050s (RCPs).

| Scenario | Radioactive forcing ($W/m^2$) | GEI concentration by the year 2100 (ppm $CO_2$ equivalent) | Temperature change (°C) by 2100 | | Greenhouse gas emissions | Agricultural Area | Publication |
|---|---|---|---|---|---|---|---|
| | | | Mean | Range | | | |
| RCP 2.6 | 2.6 | ~490 | 1.0 | 0.4–1.6 | Very low. | Medium for cropland and pasture. | (Riahi et al., 2007). |
| RCP 4.5 | 4.5 | ~650 | 1.4 | 0.9–2.0 | Medium-low mitigation (very low baseline). | Very low for both cropland and pasture. | (Fujino et al., 2006). |
| RCP 6.0 | 6.0 | ~850 | 1.3 | 0.8–1.8 | Medium baseline (high mitigation). | Medium for cropland but very low for pasture. | (Clarke et al., 2007). |
| RCP 8.5 | 8.5 | ~1370 | 2.0 | 1.4–2.6 | High baseline. | Medium for cropland but very low for pasture. | (Van Vuuren et al., 2007). |

**Table S5.** Algorithms for environmental niche modelling included in the analysis of the suitability of macadamia and the area under the curve (AUC).

| **Algorithm** | **Method** | **Description** | **AUC** |
|---|---|---|---|
| Envelope model | BIOCLIM | It computes the similarity of a location by comparing the fi (xi) at any location to a percentile distribution of the values at known locations of occurrence. | 0.74 |
| Multivariate distance | DOMAIN | It computes the Gower distance between environmental variables at any location and those at any of the known locations of occurrence. | 0.65 |
| Additive models: Generalized additive models | GAM | Semi-parametric approach to predicting non-linear responses to a suite of a predictor. | - |
| Regression: Multivariate adaptive regression splines | MARS | It is a non-parametric regression technique that automatically models non-linearity and interactions between variables. | 0.93 |
| Stepwise GAM | GAMSTEP | Builds a GAM model in a step-wise fashion. | 0.85 |
| Mahalanobis distance | MAHAL | It considers the correlations of the variables in the data set, and it is not dependent on the scale of measurements. | 1.00 |
| Maximum entropy | MAXENT | It is a machine-learning method that estimates the species distribution probability by assessing the maximum entropy distribution so that the most spread-out or closest to uniform. | 0.87 |
| Boosted regression models: Generalized boosted regression models | GBM | Based on prediction components, where each component consists of a different weighted sum of nonlinear transformations of the predictor variables. | - |
| Generalized linear models | GLM | Generalizes linear regression by allowing the linear model to be related to the response variable via a link function. | 0.88 |
| Support vector machines | SVM | Machine-learning methods that are based on classification (C-svc, nu-svc), novelty detection (one-class-svc), and regression (eps-svr, nu-svr). | 0.86 |
| Stepwise boosted regression tree models | GBMSTEP | It is a technique that aims to improve the performance of a single model by fitting many models based on stepwise selection and combining them for prediction. | - |
| Artificial neural networks | NNET | It is a machine learning approach that employs an adaptive structure, which can be trained with application data to capture complex relationships between input and out variables. | 0.77 |
| Random Forest | RF | It is a collection of tree-structured weak learners that comprised identically distributed random vectors where each tree contributes to a prediction. | 0.94 |
| Multivariate Adaptive Regression Splines | EARTH | It is a technique that builds enhanced regression models, and it involves a forward and backward automated algorithm for the selection of predictor variables. | - |
| Stepwise generalized linear models | GLMSTEP | It includes regression models in which the predictive variables are selected by an automated algorithm that involves backward elimination or forward selection. | 0.72 |
| Mixed GAM Computation Vehicle | MGCV | It provides functions for generalized additive and generalized additive mixed modelling. | 0.74 |
| Maxlike | MAXLIKE | It is a machine-learning method that estimates the species distribution probability by assessing the maximum entropy distribution so that the most spread-out or closest to uniform. | 0.77 |
| Ensemble | ENSEMBLE | This is a weighted average of all algorithms used for modelling species distribution. | 0.88 |

**Table S6.** Summary of news reports about climate change affecting macadamia production worldwide (period 2013-2019).

| Organization | Year | Country | Main report |
|---|---|---|---|
| **Helvetas** | 2018 | Nepal | Climate change is a challenge in Nepal. Current and projected increases in maximum temperatures and decreases in summer monsoon precipitation may affect macadamia tree growth and yields. |
| **Macadamia Association of Zimbabwe** | 2020 | Zimbabwe | Macadamia nut farmers experience the poorest harvest ever in 2020 owing to the side effects of climate change. Projected reduced rains and heat waves will likely affect macadamia production in the future within the country. |
| **Agricultural Research Council** | 2016 | South Africa | Changing climate is slowly altering the South Africa agricultural landscape, and farmers will need to adjust farming practices. Climate suitability studies show that the northern parts of South Africa are shifting towards the south. The Eastern Cape and KwaZulu-Natal areas could become more suitable for growing nuts, like macadamias, while Limpopo could no longer be suitable for nut production by the year 2090. |
| **Macadamia Conservation Trust** | 2017 | Australia | The MCT in Australia reports that macadamia habitat losses and fragmentation are attributed to climate change and urban footprint. New South Wales will lose about 10 % of its macadamia growing areas due to climate change. |
| **The Standard** | 2019 | Kenya | Kenyan macadamia farmers face poor crop harvests during the 2018-19 growing season due to heavy rains during the flowering phase. Interviews with some smallholder farmers reveal that climate changes will likely continue affecting macadamia yields and hence the need for sustainable technologies for adaptation. |
| **International Pacific Research Center** | 2014 | Hawai‘i | Climate change is a prominent concern among the general public in Hawai‘i and has received significant official attention within the island nation. Hamilton reports that climate change will complicate agricultural planning in Hawai‘i, especially for perennial tree crops such as macadamia and coffee. |

### Supporting R packages

Datasets were organized using the R packages “data.table” (Dowle & Srinivasan, 2020), “magrittr” (Bache & Wickham, 2016), and “tidyverse” (Hadley Wickham et al., 2019) were utilized. Layers were processed using the packages “sp” (Bivand et al., 2013), “sf” (Pebesma, 2018), “maptools” (Roger et al., 2020), “raster” (Hijmans *et al*., 2020), “rgdal” (Pebesma, 2018), “dismo” (Elith & Franklin, 2017), “rJava” (Urbanek & Urbanek, 2020) and “rgeos” (Bivand et al., 2013). Production of the maps was conducted through the usage of “ggplot2” (Wickham, 2020) and “patchwork” (Pedersen, 2020) packages.
